# Stress and behavioral correlates in the head-fixed method

**DOI:** 10.1101/2020.02.24.963371

**Authors:** Konrad Juczewski, Jonathan A. Koussa, Andrew J. Kesner, Jeong O. Lee, David M. Lovinger

**Affiliations:** Section on Synaptic Pharmacology & In Vivo Neural Function, Laboratory for Integrative Neuroscience, National Institute on Alcohol Abuse and Alcoholism, US National Institutes of Health, Rockville, Maryland, USA

## Abstract

Head-fixation of awake rodents is a method that allows for sophisticated investigation and manipulation of neural circuits *in vivo*, that would otherwise be impossible in completely freely moving animals. However, while it is known that head-fixation induces stress, its scale and habituation dynamics remain unclear. Thus, interpretation of physiological and behavioral experiments would greatly benefit from the characterization of the stress response. In our study, we used the Mobile HomeCage system (Neurotar, Finland) where animals are head-fixed to an aluminum frame, but otherwise freely moving in an ultralight carbon container floating above an air-dispensing base. To better understand this experimental environment, we analyzed locomotion and stress during an extended habituation protocol. For 25 consecutive days, mice were prepared as they would be for recording experiments, i.e. head-fixed while standing on the air-lifted platform for 2 hours per day. Throughout 25 days, blood samples were taken periodically from the tail vein to measure variation in the stress-related hormone, corticosterone. These data were compared and contrasted with behavioral data including locomotion during the 2-hours head-fixed habituation sessions and several classical behavioral measurements known to be affected by chronic mild stress.

## Introduction

Stress can have profound effects on both an animal’s behavior^1,2^ as well as physiology^3,4^. Therefore, understanding how stress might factor into awake-behaving animal experiments is important for interpreting results and making valid conclusions. This is particularly relevant when adopting new techniques, as often times the levels of stress induced by these techniques has not yet been systematically examined, and rather has been suggested only anecdotally, when compared to more established and classical behavioral paradigms. One such technique is the securing of the head of an unanesthetized animal, or head-fixing, to avoid effects of head movement during behavioral and neurophysiological experiments. Although head-fixation has been used for several decades in neurophysiological studies in monkeys^5,6^, it is only more recently that this technique has been gaining popularity in neurobiological studies using rodents, most typically mice. This renaissance was sparked by new experimental environments and modern neurobiological techniques such as optogenetics and high-resolution brain imaging^7,8^. By combining these approaches with head-fixation, investigators have been able to elucidate neural mechanism of behavioral processes at unprecedented biological resolution.

What is common for all the standard head-fixation techniques is a metal plate surgically placed on the mouse’s head such that the plate can then be attached with screws to a set of clips that restrict head-movement. However, a variety of head-fixation techniques can be distinguished based on the extent of body and head immobilization. One type of technique is to fully restrain the torso of the animal after head-fixation^9,10^. It is a simple, compact and affordable system. However, it creates a restricted environment for mice that is likely anxiogenic and reduces the behavioral repertoire available for analysis and interpretation of results. A second type of head-fixation technique involves free running on a treadmill or a spinning disc^7,11^. It allows for limb movement and some aspects of locomotor behavior but forces the animal to an unnatural body alignment and limits its possible movements to linear acceleration and deceleration. Building further is the floating-ball approach, where the mouse can move on top of a spherical ball lifted by air^12,13^. This arrangement allows a greater range of movement, but the body posture is still uncomfortable and the whole system is large and bulky. Finally, the newest approach involves a system called the mobile homecage (MHC)^14,15^, which is comprised of an air-lifted platform placed under the head-fixation frame. In contrast to other systems, it constitutes a less restricted environment giving the mouse the opportunity for body movements in multiple directions and it allows the mouse to keep more natural body posture. The MHC system is also quite compact compared to floating-ball techniques and can be easily incorporated into existing experimental set-ups and apparatuses. While the MHC system provides a more naturalistic environment that makes results more relatable to freely moving experiments, past studies using the MHC system have not provided in-depth analysis of indices of stress produced by this head-fixation technique, so the generalization of conclusions made using this technique are unclear. The present study aimed to fill this knowledge gap by systematically measuring levels of corticosterone, a stress hormone that is a common index of an animals perceived stress level^4^, and also examining stress-related behavioral phenotypes during a twenty-five-day head-fixation protocol.

## Results and discussion

### Preparation for the head-fixed experiments

We first developed a standardized method for habituating mice to head-fixation and the MHC apparatus (Fig. 1a, b), based on information from Kislin, et al. ^14^. In our experiments we wanted to find out the initial level of stress in the head-fixed situation, and if stress indices are reduced following habituation to head-fixation. All the animals (the head-fixed and the controls) used in all the protocols underwent the head-plate attachment surgery followed by two days of post-operative care. A week after the surgery every animal was habituated in two 15-minute sessions (one session a day), first to the experimenter then to the flannel that had been used to transfer animals between the home cage and the head-fixed apparatus (Fig. 1c). Finally, throughout days of the different head-fixed protocols, mice were placed in the head-fixed apparatus for 120 minutes every day with the floating container underneath. Every 5 days we took blood samples from the tail vein at the end of the head-fixed session (Fig. 1d) and measured variations in blood concentrations of the stress-related hormone, corticosterone (for review, see^4^). The control animals were only wrapped in the flannel for about a minute and left in the cage in the same room as their head-fixed littermates for the time of the head-fixed session.

**Figure 1.**
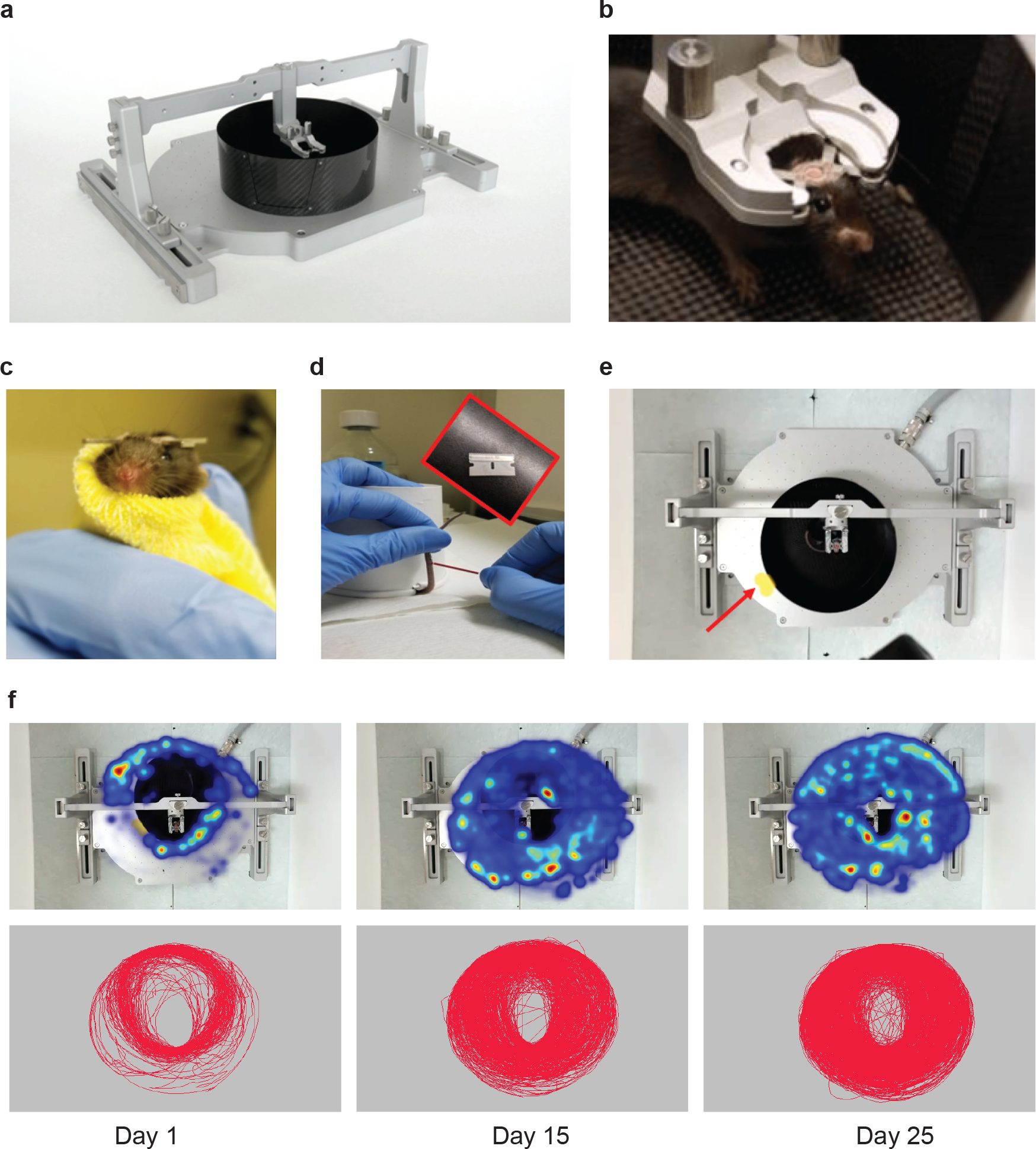
Overview of the methods. (a) Head-fixed apparatus: the air-lifted carbon fiber container (in black) placed under the head-fixation frame on the air-dispensing base. (b) Mouse with surgically-attached head-plate fixed to the head-plate holder in the head-fixation frame. (c) Mouse wrapped in the flannel for transfer between the home cage and the head-fixed apparatus. (d) Mouse partially restrained with a paper cup covering its body with a tail kept outside of the cup ready for blood sampling collection with a glass capillary after a tail-snip with a razor blade. (e) Video frame from the camera mounted on the top of the head-fixed apparatus to collect information for the locomotion analysis. Red arrow points at the yellow sticker attached to the floating container that was tracked with the EthoVision software for the movement analysis. (f) Example of heatmaps (top panel) and track maps (bottom panel) extracted with the software for locomotion analysis from the recorded videos at day 1, day 15 and day 25.

Preparatory experiments were performed on mice generated by crossing the Emx1::Cre and Ai32 reporter lines, animals that were readily available in our laboratory at the time. In the 10-day preparatory protocol the mice were head-fixed 10 times for 120 minutes/day and blood samples were collected at the end of the session on days 1, 5 and 10. We found out that the corticosterone level was reduced over the 5-day habituation and the full 10-day protocol reduced it even further (Supplementary fig. S1a). We also ran another 5-day preparatory study in which blood sampling was performed 3 times (0, 60 and 120 minutes) per every head-fixed session. In the head-fixed animals we observed a clear increase in the blood corticosterone concentration 60 minutes after the initial blood sampling through the end of the session at 120 minutes (Supplementary fig. S1b-f). In contrast, we did not see any large changes in the non-head-fixed control animals where the corticosterone level was always low. Based on these promising results from our preparatory experiments, we ran an extended 25-day habituation protocol to observe further habituation dynamics. We decided to run 120-minute head-fixed sessions using the mouse strain most commonly used in neurobiological experiments, C57Bl6/J. We also chose to collect blood samples every 5-days to avoid problems related to potential tail injuries due to frequent tail snips.

### Decreased of the head-fixation-related corticosterone changes and weight loss in the extended 25-day habituation protocol

After the preparation phase that consisted of the head-plate surgery, 7 days of recovery time and 2 days of handling, we ran the extended 25-day habituation protocol. It was followed by 4 days of behavioral testing in classical stress-associated paradigms (Fig. 2a). In the 25-day protocol we found that the initial blood corticosterone concentration after the first head-fixed session was 210.9 ± 22.46 ng/ml (Fig. 2b), a level that was about 9 times higher than the baseline level measured in the control animals, 24.2 ± 7.05 ng/ml. Corticosterone level decreased consistently over the course of the habituation protocol, but it was only after 25 head-fixed sessions that the difference between the groups was no longer statistically significant (Fig. 2b, c). In fact, there were two significant drops in the corticosterone level, between day 1 and 5 (reduction to 69 % of day 1 levels), and between day 5 and 25 (down to 35 %) (Fig. 2d). We stopped head-fixing animals on day 25, however, in 4 cases we still measured corticosterone concentration on subsequent days. It remained stable in both groups dropping insignificantly between day 25 (69.7 ± 18.91) to day 30 (32.0 ± 14.65) and staying at this level through day 60 (27.2 ± 10.82) in the head-fixed group (Fig. 2c).

**Figure 2.**
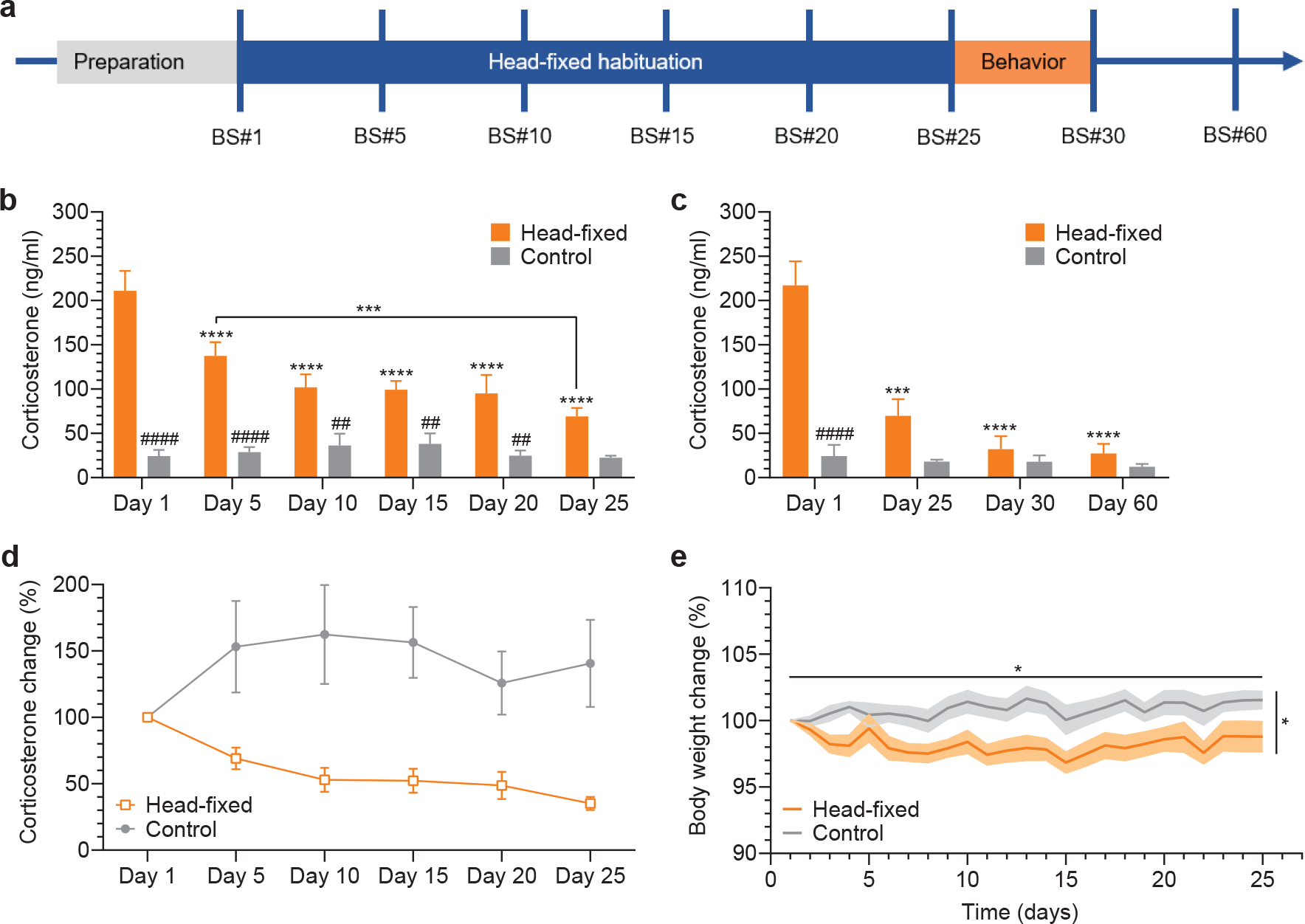
Corticosterone and body weight dynamics in the 25-day head-fixed protocol. (a) Timeline of the experiments; blood sampling (BS) every 5 days (BS#1, BS#5, etc.) at the end of the session. (b) Statistically significant difference in the blood corticosterone concentration between the groups, two-way RM ANOVA: interaction, group, and time (p < 0.0001). Sidak’s multiple comparisons between groups: difference at all days (##p < 0.008; ####p < 0.0001) except for D25 (p = 0.0755). Tukey’s multiple comparisons within groups: in the head-fixed group – D1 vs. every other day (***p = 0.0003, ****p < 0.0001) and D5 vs. D25 (**p = 0.0053) differed significantly, and in the control group – no difference in any comparison (p > 0.999); n = 8 in each group. (c) Difference in the corticosterone level between the groups statistically significant from D1 to D20 at D25 became statistically non-significant; two-way RM ANOVA: interaction (p < 0.0001), group (p = 0.0003), and time (p < 0.0001); Sidak’s multiple comparisons between groups: statistically significant difference at D1 (####p < 0.0001); no difference at D25, D30, D60, p > 0.07; Tukey’s multiple comparisons within groups: in the head-fixed group – D1 vs. D25, D30, and D60 differed significantly (****p < 0.0001), and in the control group – no difference in any comparison (p > 0.99); n = 4 in each group. (d) Percentage change in the blood corticosterone concentration normalized to D1; n = 8 in each group. (e) Changes in the body weight during the 25-day protocol; drop of weight in the head-fixed group leading to a statistically significant difference in comparison to the control group at day 5 until the end of the protocol; two-way RM ANOVA, interaction (p = 0.0756); group (*p = 0.0048); time (*p = 0.0479); Sidak’s multiple comparisons between groups: statistically significant difference in all the comparisons (p < 0.05) except for D1, D2 and D5 (p > 0.9); n = 8 in each group.

Chronic stress in rodents may result in decreased food intake leading to weight loss^3,16,17^. Therefore, we also recorded the weight of the head-fixed and control animals before every head-fixed session. In adult mice (after about 12 weeks of age) weight is stable and increases at a very slow pace^18^. This is what we saw in our control group: the average weight oscillated around 30.7 ± 0.03 g but it slightly increased from day 1 (30.43 ± 0.63 g) to day 25 (30.91 ± 0.73 g) of the studies. In contrast, in the head-fixed group, the average weight oscillated around 29.5 ± 0.04 g but it decreased with time from day 1 (30.0 ± 0.60 g) to day 25 (29.6 ± 0.62 g). In fact, in the head-fixed group weight dropped after the first head-fixed session leading to a statistically significant difference between the groups already at day 3 (98.2 ± 0.65 % versus 100.5 ± 0.5 % of the initial body weight, the head-fixed and the control group, respectively). The difference in weight between the groups, despite some minor fluctuations, remained stable till the end of the 25-day protocol (Fig. 2e).

Secretion of corticosterone is influenced by various external and internal factors; therefore, we conducted control experiments and additional analysis. Firstly, we checked whether the blood sampling procedure by itself constituted a confounding factor in our experiments due to shifting the baseline corticosterone level. We ran 3 blood sampling control experiments where animals underwent the same initial procedures as the animals in the 25-day protocol, but they were withdrawn from the study after a single blood sampling at day 5, 15 or 25. In the head-fixed group we did not observe any major difference in the corticosterone level between those animals and the corresponding animals that were blood-sampled several times during the 25-day protocol (Supplementary fig. S1g). However, in the control group the similar comparison showed a higher level of corticosterone at day 5 and 15 in the animals that were blood-sampled several times (Supplementary fig. S1h). Moreover, knowing that in rodents the corticosterone level follows a circadian rhythm^19,20^, we counter-balanced our experimental groups in the 25-day protocol. We chose four time points for the head-fixed sessions and we looked at the blood sampling results in that context. We saw the expected increase in the blood corticosterone concentration throughout the day, with major differences between the morning- and the late afternoon-sampled animals (Supplementary fig. S1i). In contrast, in the head-fixed group this pattern was interrupted. To sum up, control experiments/analysis showed that less frequent blood sampling (every 5 days) and counter-balancing the groups in terms of the time of the experiment were important for an optimal experimental design.

### Repeated head-fixation does not alter several stress-sensitive behavioral phenotypes but results in altered fluid intake

We next examined if the 25-day fixation protocol produced any long-term behavioral effects. This information may be especially useful in the context of learning and plasticity that often require complex training for several days^21,22^. We used a few classical behavioral measurements known to be affected by chronic mild stress^3,23,24^. We started on day 26 in the morning with the open field test that is used to analyze general locomotor behavior but also the anxiety/stress related to open space. We did not see any difference between the head-fixed and the control group in the total distance traveled (Fig. 3a). Also, the time spent in the center of the field was similar between the groups as well as the number of returns to the central part of the field and the latency to the first visit in the center (Fig. 3b; Supplementary fig. S2a, b). Six hours later, in the afternoon, we performed a forced-swim test that is used to assess instinctive survival behavior in extreme circumstances that may be affected by stress. We did not find any statistically significant change in the measured parameters: latency to the first floating event, total floating time, the number of floating events and the number of feces left in the water (Fig. 3c, d; Supplementary fig. S2c, d).

**Figure 3.**
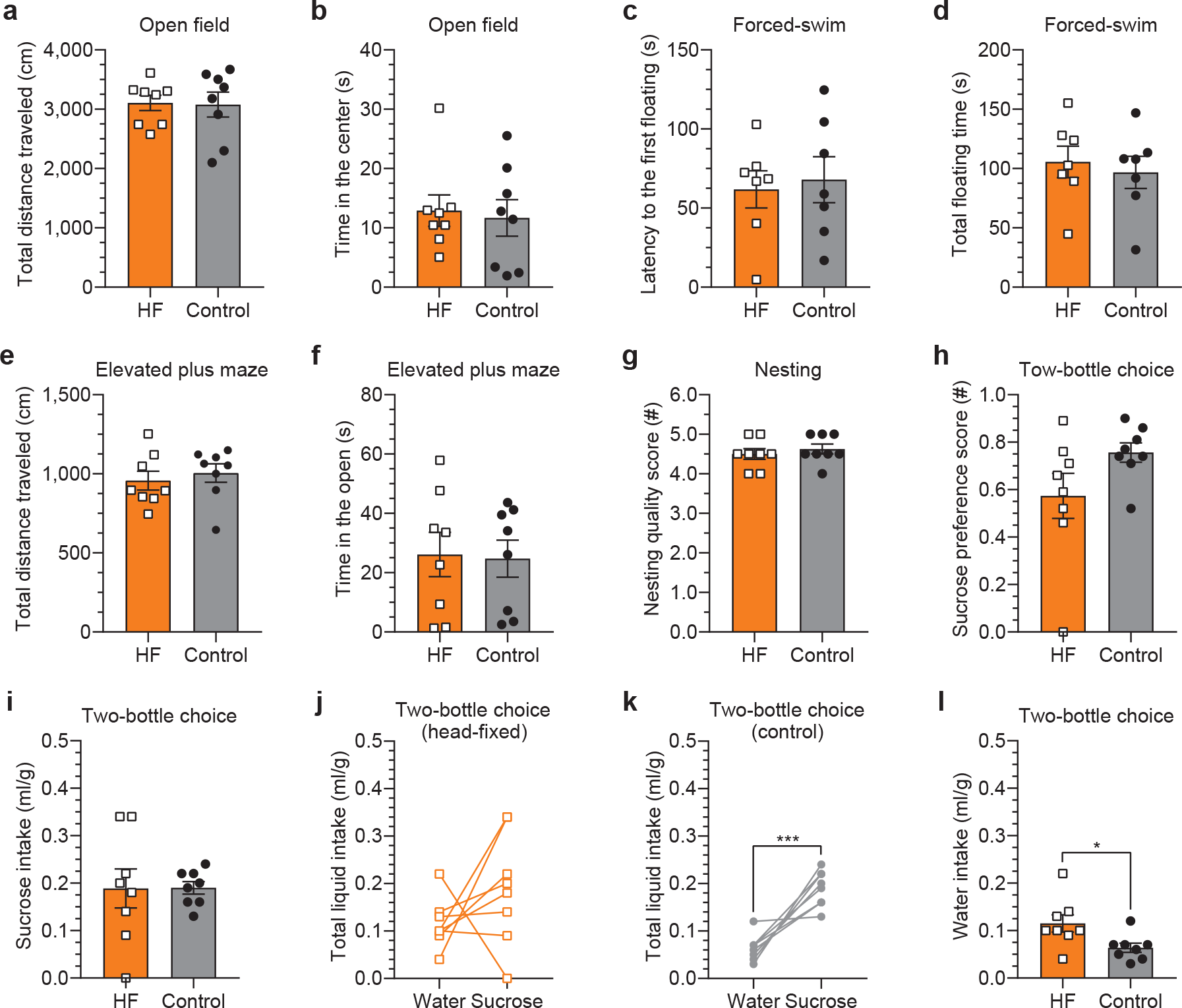
Long-term effects of the 25-day head-fixation tested in the stress-associated behavioral paradigms. (a, b) Both groups traveled similar distances during the trial and spent similar time in the center of the open field box; p = 0.9238 and p = 0.6085, respectively. (c, d) Both groups had similar latency to the first floating event and spent similar time floating in the forced-swim test; p = 0.7342 and p = 0.6770, respectively. (e, f) Both groups traveled similar distances during the trial and spent similar time in the open arms of the elevated plus maze; p = 0.6711 and p = 0.9163, respectively. (g) Both groups built nests of similar quality using all the available nesting material; scale 1 to 5 (worse to the best quality score, respectively); p = 0.6263. (h-l) Two-bottle free choice task used to test sucrose preference; liquid consumption adjusted for the body weight. (h) No change in the sucrose preference score in the head-fixed group relative to controls; scale 0 to 1, 0 = 100% water and 1 = 100% sucrose preference; p = 0.7563 (i) No difference between the groups in the total volume of consumed water (p = 0.8390). (j, k) Lack of statistically significant difference between the total volume of water versus sucrose consumed in the head-fixed group (p = 0.2400) and a significant difference in the control group (***p = 0.0005). (l) Statistically significant difference between the groups in the total volume of consumed water (*p = 0.0435); n = 8 in each group.

In the morning of the second day of behavioral experiments we performed an elevated plus maze test. This task relies upon rodents’ proclivity for dark, enclosed spaces by measuring their preference to spend time in the open or closed arms of the maze. Also, in this task the head-fixed and control groups did not differ on any of the measured parameters (Fig. 3e, f and Supplementary fig. S2e-l). The mice traveled similar distances during the time of the trial. They spent similar short times in the open arms of the maze, preferring the closed arms. In the late afternoon of the same day, just before the dark cycle, we moved all the mice to individual cages with 3 grams of nesting material in each cage to measure the quality of their nests. We scored the nest quality on the next day using the scale standardized by Deacon ^25^. This task is thought to provide an index of apathy/anhedonia which may be reflected by a lower quality of nest construction. We did not observe any group difference in nest quality. Both groups shredded the majority of the provided nesting material similarly and assembled it into a good quality nest, with the average score of 4.5 out of 5.0 points (Fig. 3g).

Another known measure of apathic/anhedonic behavior is sucrose preference tested in the two-bottle free choice task. Thus, on days 3 and 4 of behavioral testing, we ran the two-bottle free choice task where one bottle was filled with 1-%-sucrose and another with regular drinking water. In regular conditions mice, just as humans, prefer sweet liquids over water but they do it for a higher caloric content not for its sweet taste^26^. While there was no inter-group difference in sucrose preference (Fig. 3h), there was considerable variability in sucrose intake and preference in the head-fixed group as compared to the control group (Fig. 3i-k). In addition, the head-fixed animals consumed significantly more water than the controls (Fig. 3l). These results together suggest a change in fluid intake that might relate to excess thirst in the head-fixed mice even a few days after the last head-fixed session.

### Locomotion pattern does not predict changes in the head-fixed stress, but distance traveled is correlated with corticosterone level

In both humans and rodents physical exercise, such as voluntary wheel running, promotes stress resistance, meaning a reduced response to stressor exposure^27,28^. Because head-fixed mice in the MHC can easily move the floating container with their paws, we collected locomotion data to see whether movement in this setting is related to measures of the blood corticosterone concentration. We performed video tracking of a yellow sticker attached to the MHC container which provided a reliable report of the mouse’s body movement (Fig. 1e). Examination of movement-related heatmaps and track maps gave us a general idea about the mouse’s locomotion (Fig. 1f). Next, we characterized it further by quantifying movement time, average velocity and total distance traveled during every session throughout the 25 days of training. Mice exhibited greater locomotion on the first day (24.61 ± 4.27 % of the session time) relative to subsequent days (Fig. 4a). The average movement time then significantly dropped to 14.2 ± 2.78 % on day 2. Finally, it stabilized at about 20 % on day 5 remaining at this level with some minor fluctuations till day 25 (average movement time over 25 days was 20.5 ± 0.52 %). In contrast, total distance traveled as well as maximum velocity per day consistently increased across habituation days after day 1 (Fig. 4b, c).

**Figure 4.**
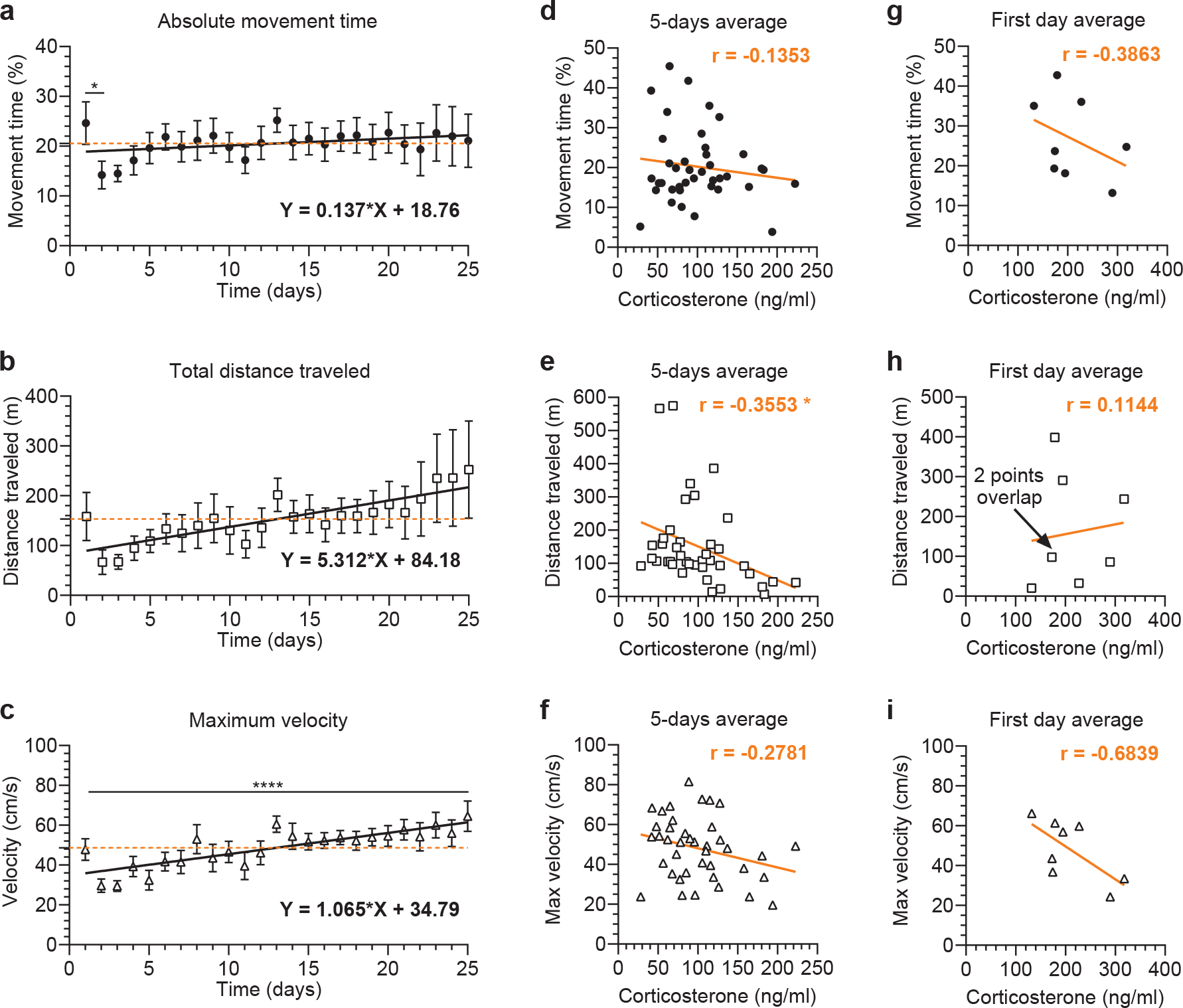
Locomotion dynamics and correspondence to the overall corticosterone level in the head-fixed animals. (a-c) Locomotion parameters in the 25-day protocol; daily averages were calculated with a resolution of seconds; the dashed orange line corresponds to the grand average and the solid line to the linear regression. (a) Movement time stabilized at about 20 % after 5 days of the head-fixed protocol, but all fluctuations except for the drop between D1 and D2 were not statistically significant, one-way RM ANOVA (p = 0.3607). Dunnett’s multiple comparisons test for D1 vs. D2 (*p = 0.0396) and for D1 vs. all other days (p > 0.3). (b, c) No statistically significant change in the total distance traveled per day and significant increase in the daily maximum velocity with the training time, one-way RM ANOVA (p = 0.3607 and ****p < 0.0001, respectively). (d-f) Correlation between corticosterone blood concentration measured every 5 days and locomotion parameters calculated as averages for the time in-between blood sampling during the 25-day protocol (averages for day 2 to day 5, day 6 to day 10, etc.). (d) No statistically significant correlation between blood corticosterone and the absolute movement time (r = –0.1353, p = 0.4051). (e) Statistically significant correlation between blood corticosterone and the total distance traveled (r = –0.3553, *p = 0.0244). (f) No statistically significant correlation between blood corticosterone and the maximum velocity (r = –0.2781, p = 0.0823). (g-i) Correlation between the blood corticosterone concentration and the locomotion parameters from the first day (omitted in the 5-day analysis above); no statistically significant correlation between the corticosterone and the absolute movement time, the total distance traveled and the maximum velocity (r = –0.3863, p = 0.3446; r = 0.1144, p = 0.7873; r = –0.6839, p = 0.0614, respectively); the solid black or orange line represents linear regression; all comparisons made with Pearson correlation analysis; n = 8.

Knowing corticosterone dynamics and changes in the movement parameters throughout the 25-day protocol, we checked whether any of these changes were related to decreasing corticosterone level. We did not find any significant correlation between the movement time or average velocity and the corticosterone level (Fig. 4d, f). However, there was a significant negative correlation between the distance traveled and the corticosterone level (Fig. 4e). The longer distance the animals ran, the lower the level of corticosterone. Furthermore, we also tested whether any of the locomotion parameters measured on the first day can be used as a predictor of the day 1 corticosterone level. We did not see any statistically significant correlation with the movement time, the average velocity or the distance traveled (Fig. 4g-i). These results suggest that the locomotion pattern cannot be used as a reliable biomarker of the stress level in the head-fixed mice.

### Dynamics of the voluntary locomotion behavior in the head-fixed mouse

In the head-fixed experiments mice and rats are often trained to perform complex sensory-motor tasks while their brain activity is being monitored with sensitive recording techniques such as patch-clamp electrophysiology^8,29^. Spontaneous voluntary movement, i.e. running in the MHC, may affect training efficiency as well as hinder neural recordings or interpretation of the results. Thus, determination of the optimal experimental design for understanding locomotion dynamics will be useful for experimenters, especially if they are interested in studying/avoiding voluntary movement in this head-fixed setting. We found that mice were showing relatively short bursts of movement spread out over much of the duration of the session which was clear from the movement time analysis. The absolute movement time calculated on a per frame scale (the frame rate = 30 Hz) showed that, on average, the mice were moving 5.5 ± 0.32 % of the 120-minute session (Fig. 5a). In the same time, if summing up every minute when some movement occurred, it was about 60 to 90 minutes every session (Fig. 5b). Hence, the total distance traveled had been increasing constantly from the first to the last minute of the session (Fig. 5c). To learn whether there was any general trend when the movement occurred, we analyzed the first and the second hour of every session. We averaged all days for each animal separately taking into account individual differences. It was clearly visible that most of the animals were more active and traveled longer distances during the first hour (Fig. 5d, e). However, this difference was not to statistically significant in the grand average summaries (all animals all days together) due to high individual variability between the animals (Fig. 5f, g).

**Figure 5.**
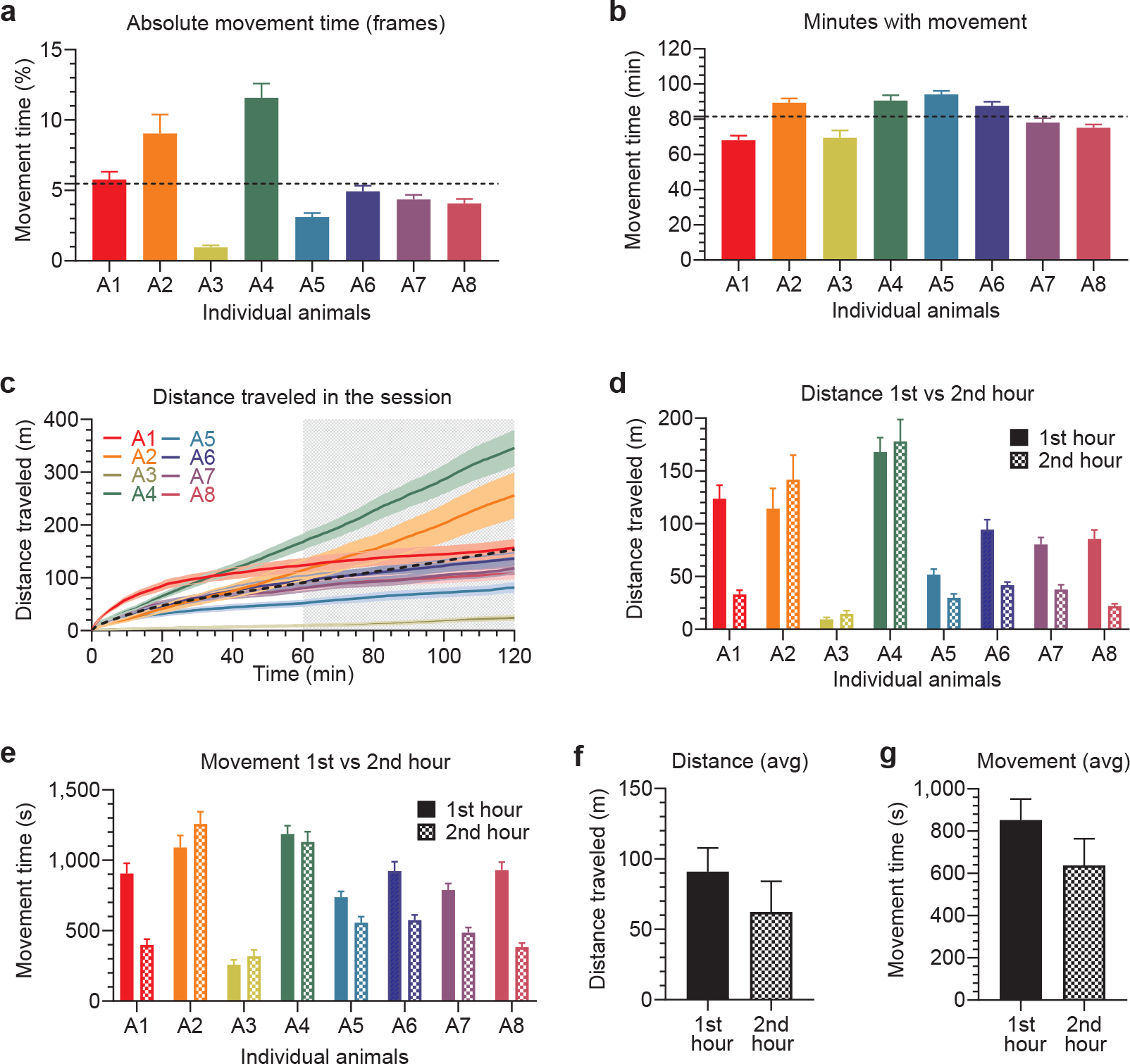
Locomotion dynamics during the 120-minute head-fixed session. (a-f) Movement and distance analysis, averages from all the 25 head-fixed sessions. (a, b) Individual differences between the head-fixed animals in the absolute movement time (proportion of frames with movement recorded at 30 Hz resolution) and in the minutes with movement per head-fixed session (when counting every minute with any movement), respectively; dashed line represents the grand average from all the sessions and all the animals. (c) Average distance during every minute of the 120-minute head-fixed session, all days together; dashed line corresponds to the grand average from all the sessions and all the animals together. (d-g) Movement time and the total distance traveled during the first and the second hour of the head-fixed session calculated with a resolution of seconds. (d, e) Individual animal data for the movement time and for the total distance traveled. (f, g) No statistically significant change in the average movement time and total distance traveled between the first and the second hour of the head-fixed session (paired t-test, p = 0.0514 and p = 0.0882, respectively); n = 8.

### Individual differences and motor skill refinement in the floating container

Moving the air-lifted floating container in a controlled manner demands practice that leads to the improvement in speed, accuracy and consistency of movement with training. Hence, it can be classified as a motor skill^28^. In our study, refinement of this skill happened at a different pace for different animals. Therefore, we also quantified movement time of individual animals throughout 25 days of habituation. We found 3 types of animals: animals that increased, decreased or always kept movement engagement at the same level (Supplementary fig. S3a-c, respectively). Nevertheless, all of them seemed to develop a better control over the floating container with time. Thus, in the next step we characterized the movement quality by focusing on changes in velocity in every session.

We knew from our observation that the majority of mice in the first session/s did not know how to operate on the floating container and their movements seemed to be uncontrolled. With training time, initial uncertain and random movements were replaced by organized exploratory behavior and smooth-running events, a sign of a more controlled movement technique. To quantify this transition, from velocity analysis we extracted *bouts of activity* that were defined as temporary changes in velocity as opposed to the resting time when a mouse did not move the floating container (Supplementary fig. S4a-c). We focused on two aspects that may be helpful in describing movement control, namely duration and velocity of individual bouts. Improvement in movement control, was measured as significant increases in the average velocity of a bout while there were no significant changes in the average duration of a bout throughout the head-fixed protocol (Fig. 6a, b). To characterize movements in detail, we also classified the bouts into 4 categories by their duration, short and long, and by their velocity, slow and fast (for details see Methods and Supplementary fig. S4d). Interestingly, there was a similar proportion of short and long bouts throughout the 25-day protocol (on average 48.5 ± 0.92 % and 51.5 ± 0.92 %, respectively). However, what changed the most was the increase in the number of fast bouts, both long and short (9.9 ± 3.69 % to 28.2 ± 6.39 % and 0.9 ± 0.34 % to 4.6 ± 1.26 %, respectively) at the expanse of long and slow bouts (43.4 ± 4.22 % and 21.3 ± 3.98 %) (Fig. 6c). The increase in fast bout proportion corresponded to our observation that with time mouse movements became smoother and the animals were able to speed up in both respects, the exploration and the running. Finally, knowing changes in the corticosterone level throughout the 25-day protocol, we were curious whether any of the aforementioned bout types are related to the corticosterone drop. Indeed, we found a significant positive correlation between long and slow bouts and a decrease in the corticosterone level but no significant correlations with other bouts (Fig. 6d-g). Because bout dynamics may be affected by training time, we also examined the relationship between training day and bout proportion. We found that changes in the proportion of the long bouts (both fast and slow) over time were significant whereas changes in the proportion of the short bouts were not (Fig. 6h). Therefore, decreased proportion of the long and slow bouts is related to the lower corticosterone level, not only a reflection of the training time.

**Figure 6.**
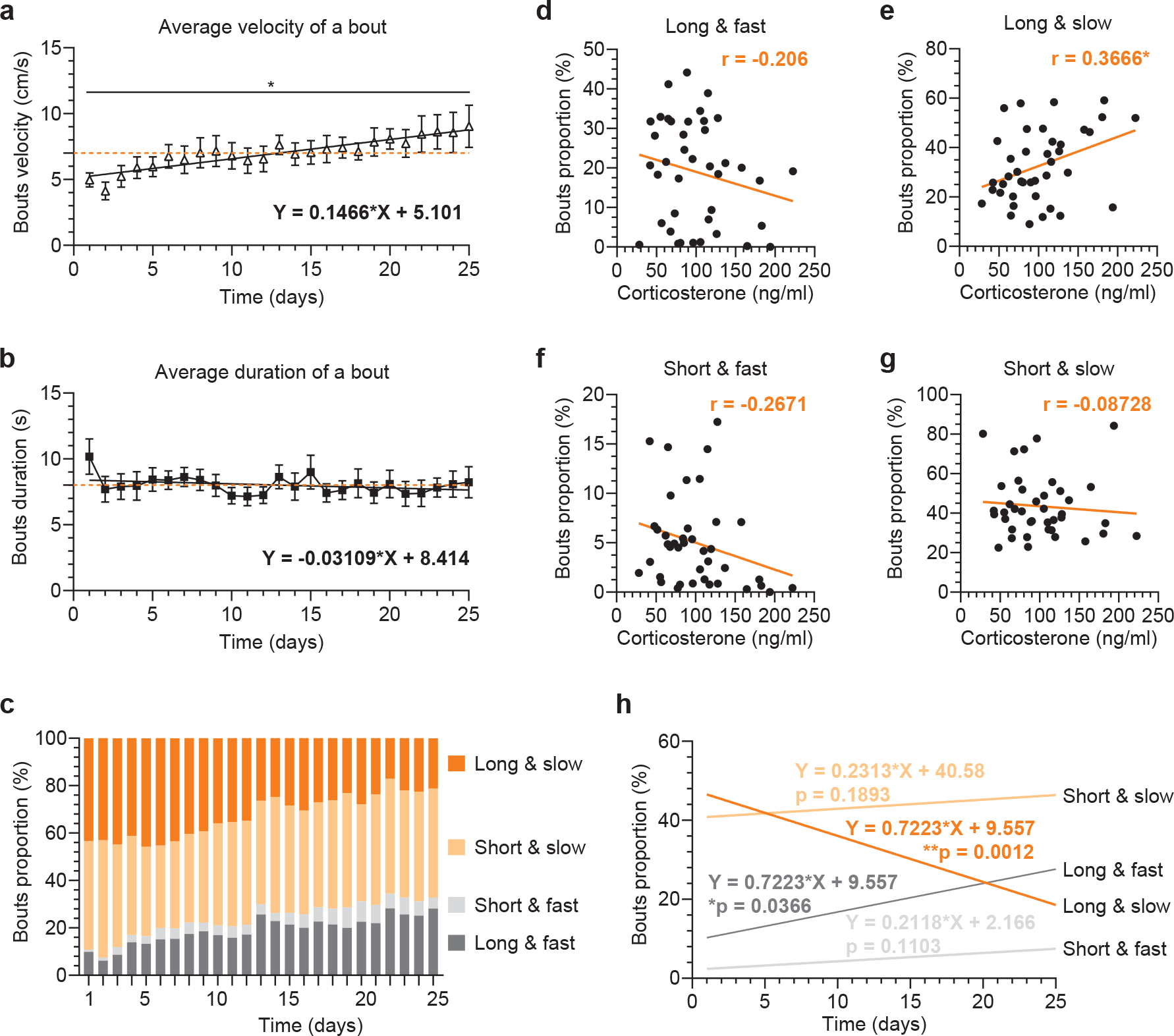
Motor control measure with the container movement and correlation with lower levels of blood corticosterone. (a-c) Detailed velocity analysis based on the *bouts of activity* in the 25-day protocol (for details see Methods); the dashed orange line corresponds to the grand average and the solid line to the linear regression. (a) Statistically significant increase in the average velocity of a bout with passing time, one-way RM ANOVA (*p = 0.0307). (b) Stable average duration of a bout throughout the 25-day protocol, no statistically significant changes, one-way ANOVA (p = 0.2293). (c) Bouts of activity dynamics, changes in the proportion between 4 different categories of bouts with passing time. (d-g) Correlations between the blood corticosterone concentration measured every 5 days and changes in bout proportion. Measures were calculated as averages for the time in-between blood sampling during the 25-day protocol (averages for day 2 to day 5, day 6 to day 10, etc.). Significant correlation between the corticosterone level and the long & slow bouts (r = 0.3666, *p = 0.0200) but no statistically significant correlation with other types of bouts: long & fast, short & fast and short & slow (r = –0.206, p = 0.2021; r = –0.2671, p = 0.0957; r = –0.08728, p = 0.5923; r = 0.3666; respectively). (h) Statistically significant changes in proportion of the long bouts with the training time and no changes in the short bouts, one-way ANOVA, long & fast (*p = 0.0366), long & slow (**p = 0.0012), short & fast (p = 0.1103), short & slow (p = 0.1893); n = 8.

### Detailed analysis of container-spinning behavior and other control experiments

In our study, the majority of mice began to *spin* the floating container as training progressed. During the early training sessions, the container was moved in all directions but with time the mice gradually shifted to one side of the container and spent more time running in proximity of the walls, moving the container in a single direction (a movement that resulted in spinning of the container). We could not easily quantify this movement by tracking the yellow sticker attached to the container. Thus, we used the MHC Tracking System (Neurotar Ltd, Finland), a special system based on magnetic sensors designed for detailed locomotion analysis in the MHC set up (for details see Methods). In the 5-day protocol of our supplementary study (5 head-fixed sessions, 60-minute each), we studied spinning behavior. It was clear from the color-coded tracking maps that mice gradually developed controlled spinning behavior (Fig. 7a). They spent more time closer to the container walls as the time passed, within the session and across training days. Furthermore, we extracted place preference data dividing the container into two zones, the walls and the middle, to quantify the proximity to the wall (Fig. 7b). Indeed, mice spent more time in the wall zone than in the middle zone once they gained better control over the spinning movement.

**Figure 7.**
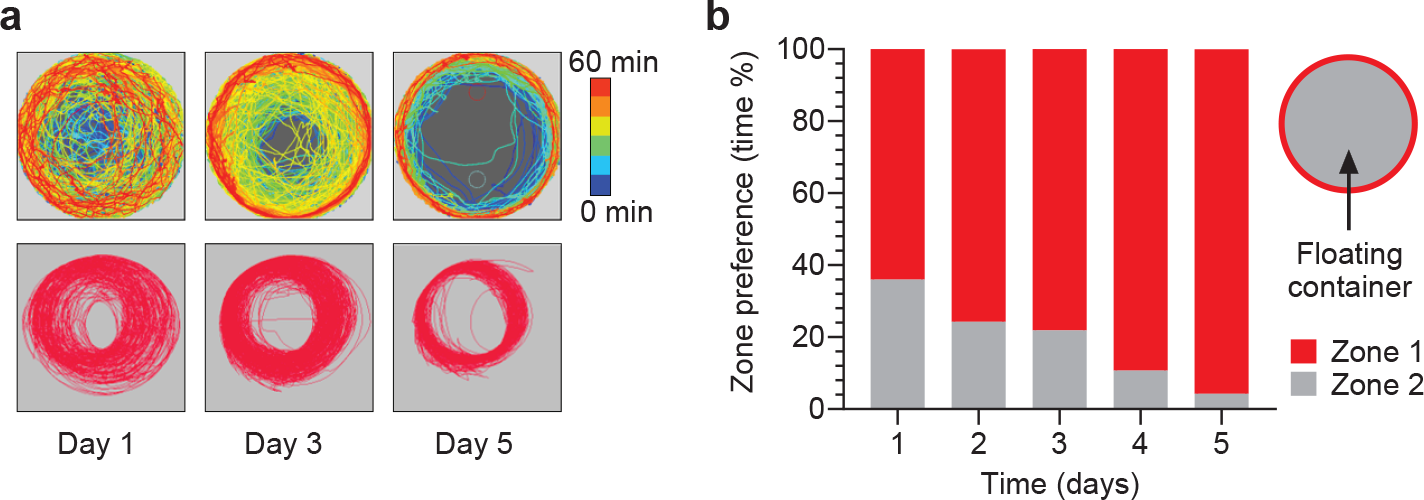
Spinning behavior analysis in the 5-day head-fixed protocol. (a) Track maps collected with the MHC Tracking System (top panels) and with the EthoVision software (bottom panels). In the MHC Tracking System, the additional option of colorometric presentation was available (from blue colors to red colors reflecting the passing time of the session). Visible difference between two methods (in the MHC Tracking System analysis lines represent movement of the container relative to the mouse, in the EthoVision software analysis –movement of the yellow sticker attached to the container). Analysis of spinning behavior in the context of a preferred placement of the container was possible only with the MHC Tracking System. Mice shifted the container from the middle to the side throughout the session and ran in proximity to the walls once their control over the container was refined. (b) Zone preference data from all training days (D1-5) recorded from 4 animals presented as the average percentage of the time spend in the zone during the 60-minute head-fixed session. Schema of the zone division of the floating container (red corresponds to the walls and grey to the middle part of the container).

In the 5-day protocol, we ran two groups of animals, the *floating* group (the container was air-lifted and floating) and the *fixed* group (the container was fixed with a tape to the base). Access to voluntary running in the floating group did not affect the corticosterone levels when compared to the fixed group as concluded from the blood samples taken after the first and the last head-fixed session (Supplementary fig. S5a-c). Also, it seems that 60-minute sessions were too short to reduce the corticosterone levels in head-fixed mice over 5 days. The floating group on average ran for 1217.0 ± 76.66 seconds out of 3600 seconds in each session which was significantly longer than the struggling body movements performed by the fixed group, on average for 559.3 ± 26.1 seconds (Supplementary fig. S5d). However, in both groups the proportion of the movement time did not significantly differ across training days (Supplementary fig. S5e). We also conducted the same stress-related behavioral tests used in the 25-day protocol, but they did not reveal any statistically significant differences between the floating and the fixed groups except for two minor parameters (Supplementary fig. S5f-q). In the open field test the animals from the fixed group showed longer latency to visit the center of the box (Supplementary fig. S5h). In addition, they presented significantly lower number of attempts to the open arm in the elevated plus maze (Supplementary fig. S5n). Both changes may reflect slight anxiety in the fixed group. Interestingly, in contrast to the 25-day protocol, the floating group did not display any change in the sucrose preference test which convinced us that the fluid intake changes in the 25-day protocol may result from factors other than stress.

## Conclusions

Measurement of blood corticosterone concentration is a commonly used method to assess physiological stress response^30–32^. However, it is important to remember that stress is a complex physiological mechanism incorporating changes in secretion of many different hormones and transmitters, in addition to corticosterone (for review, see^4^). Therefore, any measurements of the blood corticosterone concentration related to particular stressors should be considered as an indicator of potential stress, not an ultimate stress index. This is especially important since the corticosterone level oscillates cyclically following the circadian rhythm in males and the estrus cycle in females^19,33^. In our studies we counter-balanced groups for the circadian factor to be certain that what we recorded was related to head-fixation, not a natural circadian change in the corticosterone level. Furthermore, we used only males to avoid fluctuations related to the estrous cycle^19,33^ which would be especially difficult to overcome in the extended 25-day head-fixed protocol. Our head-fixed experiments revealed that the initial level of corticosterone in the blood sample after a single head-fixed session is almost 2-6 times lower^34^ in the MHC approach than in the full-body restraint, about 200 ng/ml versus about 400-1200 ng/ml depending on the source of the ELISA assay^30^, the basal level of the corticosterone^35^ and the study design^19,36,37^. However, this value is still 9 times higher than the corticosterone level in the non-head-fixed control animals (about 20 ng/ml) which is lower or corresponds to the baseline corticosterone level observed in other studies^19,31,34–37^. Therefore, a habituation protocol where mice adapt to the head-fixing constitutes an important experimental procedure, especially for experiments where stress is considered a confounding factor. After careful analysis of our corticosterone data we recommend a 10-day habituation protocol as a good solution if your research plan does not allow for the full 25-day or longer habituation protocol. At day 10 corticosterone level as well as mouse body weight stabilizes which is a good sign not only of reduced stress level but also adaptation to the new environment.

We did not find any statistically significant difference in stress-sensitive behaviors after the 25-day protocol apart from a change in water intake revealed in the sucrose preference test. Neither did we see any difference after the shorter 5-day protocol. Both results were similar to a behavioral study performed on animals after full-body restraint^37^. We also studied voluntary locomotion during the head-fixed protocol. Studying this simple and spontaneous behavior was important because increased stress level might affect learning of sensory-motor tasks as well as interfere with physiological recordings during their performance. We showed that mice were generally active throughout 120-minute sessions with slightly higher engagement in locomotion during the first hour. We also showed strong individual differences between animals, suggesting 3 different types of mouse behavior: movement increase, decrease or no change. It may be interesting to follow up this distinction and test whether this movement feature will correspond to other characteristics, such as learning capability or susceptibility to behaviors related to drug use disorders.

We also examined the correlation between locomotion and stress indices. We found that corticosterone levels were generally lower with the larger numbers of rotations of the MHC container. Furthermore, we also quantified movement refinement in the bouts of activity analysis in which uncontrolled movements were replaced by fully executed control over the container. This shift was reflected in the proportional change between different bout categories, namely, increase in the faster bout proportion. Nevertheless, it was only the change in the long and slow bout proportion that was significantly correlated with the level of corticosterone. Hence, the long and the slow bouts may be useful as a biomarker for the stress level assessment. Additionally, classification of movement by velocity (bouts of activity) may be a useful measurement for research dealing with the motor skill learning/refinement.

To sum up, we showed that head-fixation is associated with at least one physiological sign of stress, namely increased blood corticosterone. Therefore, the stress-sensitivity of assays must be considered when designing experiments involving head-fixation. Nevertheless, the head-fixing procedure did not strongly disrupt the refinement of movement or other behaviors neither in the longer head-fixed protocol (25-day) nor in the shorter one (5-day). Hence, the full 25-day habituation protocol does not have to be the best option for every type of experiment and shorter habituation protocols run for 5 or 10 days may be a reasonable choice.

## Methods

### Animals

All animal protocols were approved by the US National Institute on Alcohol Abuse and Alcoholism (NIAAA). All experiments were performed in accordance with the National Institutes of Health (NIH) guidelines for animal research. All mice were purchased from the Jackson Laboratory (Bar Harbor, ME, US). Only adult, male mice were used in the matching age range, 13 weeks at the time of surgery except for some animals used in the pilot experiments (8 weeks) and the supplementary experiments (20, 21 and 24 weeks). This age-range was chosen because the younger, pre-adolescent animals present higher and more variable response to stressors^38,39^. C57BL6/J animals were used in the main project and supplementary experiments and Emx1∷Cre animals crossed with the Ai32 reporter line were used in the pilot experiments. All mice were housed on a 12-hour light cycle (6:30-18:30), initially, 4 per cage. Immediately after the head-plate surgery, mice were separated and kept 2 or 1 per cage depending on the experimental group, except for the preparatory study (5-day and 10-day preparatory protocol) mice that were always kept 2-4 per cage. Access to rodent chow and water was provided *ad libitum.* 80 mice underwent the head-plate surgery of which 3 died during the surgery procedure and 10 were removed from the study due to a head-plate loss during the experimental procedures. All behavioral experiments were performed during the light phase with control and head-fixed mice run on the same tasks during the same time period.

### Head-plate surgery

Head-plate surgery was performed a week before the beginning of head-fixed experiments. Animals were placed in the stereotaxic frame under general isoflurane anesthesia (0.5-1.5 % v/v isoflurane in oxygen). The skull was leveled by adjusting ear bar and nose bar position. The skull surface between the bregma and the lambda (cranial suture points) was exposed by removing a small piece of skin above this area with scissors and forceps. Next, the exposed skull surface was cleaned with 0.1 M phosphate buffer, then 70 % EtOH and 3% Hydrogen peroxide and carefully dried with a Kim wipe. Head-plate model 8 or 9 that weighs about 1 g (Neurotar Ltd, Finland) was initially glued with superglue (World Precision Instruments, FL, US) and secured to the skull with 2 miniature screws (Antrin Miniature Specialties Inc., US) were fitted diagonally at a small angle next to the head-plate, so their heads stabilized the head-plate position. More glue was applied to a contact place between the head-plate and the skull. The connection was supported even further with dental cement (Coralite Dental Products Inc., US). At the end of surgery animals were placed in new cages in pairs and a post-operative mix was applied daily for the following two days, Lactated Ringer’s Injection, USP (Hospira, Inc., IL, US) at a dose 1 ml per 30.0 g of body weight.

### Head-fixed experiments

Mice were handled for two days in 15-minute sessions per day. On the first day they could move around on the experimenter’s hands to get accustomed to the experimenter. On the second day they were exploring small pieces of cotton flannel of a similar white/light color to avoid potential color-bias (mice are able to see colors but with inability to distinguish some colors). Flannel pieces were assigned to individual mice to prevent another animal’s smell influencing the results. Mice were then wrapped and unwrapped in the flannel pieces several times during the handling session for proper habituation, as shown with the instruction videos available online (Neurotar Ltd, Finland). After 2 days of handling (one without the flannel and one with the flannel), the first day of the head-fixed protocol started. Each animal was weighed and then wrapped in the flannel. An animal from the head-fixed group was positioned in the head-fixed apparatus. The control animal was held in the flannel for a corresponding time then it was returned to its cage which was kept in the same room as their head-fixed littermate, so that they be always exposed to the same environment. The head-fixed apparatus was the Mobile HomeCage^®^ system (Neurotar, Finland) referred to as the *MHC*, that can also be built^40^. It is a research device where animals are head-fixed to an aluminum frame, but otherwise freely moving in an ultralight carbon fiber container floating above an air-dispensing base. The circular floating container used in this study with dimensions of 325 × 70 mm weighed about 50 g. Floating the container above the air-dispensing base was accomplished using air pressure from a connected pump (Matala Water Technology Co., Ltd., US) adjusted to about 100-110 liters/minute at 2.9 psi.

Once the mouse was positioned under the aluminum frame, the head-plate was clamped in the head-post, the floating container was placed underneath, the flannel wrapping was removed, and the air-pump was turned on. The air-pump generated noise of about 70 dB and the MHC stage lighting was about 200 lux. The MHC stage was surrounded by non-transparent, white plastic walls that separated it from the rest of the room. There were several types of the head-fixed protocols used in these studies including 5-day and 10-day pilot experiments, 25-day protocol in the main study and 5-day protocol in the supplementary study. In all cases the head-fixed sessions were performed on consecutive days and lasted for 120 minutes except for the 5-day protocol in which it lasted 60-minutes. In all the pilot studies and 25-day protocol two groups were compared. The head-fixed and the non-head-fixed animals underwent similar procedures including the head-plate surgery. In the 5-day protocol all the animals were head-fixed, but the comparison was between the *floating* group (the container was air-lifted and floating) and the *fixed* group (the container was fixed to the base). Furthermore, in the floating group the MHC Tracking System (Neurotar, Finland) was used together with magnetic tracking mats inserted in the container increasing its weight to about 65 g. The air-pump pressure had to be adjusted to about 120 liters/minute at 2.9 psi to make the container float in this condition. The container should be lifted by the air to about 0.1 mm above the base to reduce friction and allow for movement. If the air-pressure is too high, the mouse cannot control the movement of the container and the bottom part of the body constantly moves sideways. To standardize air-pressure between experiments a sheet of paper should just fit in-between the base and the container with no extra space. In the fixed group we used the same container with the inserted magnetic mat. The head-fixed apparatus, a head-fixed mouse and a flannel-wrapped mouse are shown in Fig. 1a-c.

### Corticosterone blood concentration measurement

After careful consideration of different blood sampling methods (for review, see ^31,32^), tail vein bleeding from the awake animal was chosen. It was the least stressful procedure which allowed us to collect relatively large blood samples in a short time. It was easy to repeat several times in a consistent manner and helped to avoid the confounding effect of anesthesia. Blood sampling was performed immediately after the head-fixed session and each blood sample was about 40-50 ul. Mice were placed on the table under a small paper cup covering its body with a tail kept outside the cup stretched with a thumb and an index finger (Fig. 1d). The tail was snipped with a razor blade; a sample was collected with a heparin-coated glass capillary tube (Thermo Fisher Scientific, MA, US) that was closed with the Surgipath Critocap (Leica Biosystems Richmond, Inc., IL, US) at the end (Supplementary fig. 1d). The vein was secured afterward by applying gentle pressure with fingers above the cut while touching it with the Flexible caustic applicator 6” coated with silver nitrate (75 %) and potassium nitrate (25%) (Bray Group Ltd., England). The entire blood sampling procedure took no longer than 5 minutes (most often less than 2-3 minutes) minimizing confounding stress-response related to the blood sampling itself. It was reported that the corticosterone level increase in response to the stressor is noticeable no earlier than 5 minutes after its occurrence^31^. The sample was taken first from the control animal that had remained alone in the cage, then from the head-fixed littermate which was just coming back to the cage to avoid potential acute stress related to a bleeding littermate. The capillary with a blood sample was placed in the DM1424 Hematocrit Centrifuge (SCILOGEX, LLC, CT, US) and spun at 15,000g for 5 minutes at room temperature to separate the components of blood: red blood cells, platelets and plasma. About 15-20 ul of plasma that precipitated as a transparent uncolored layer was transferred to an Eppendorf tube and placed at − 20 C freezer for storage up to 4 weeks before further processing.

Blood corticosterone concentration was measured with enzyme-linked immunosorbent assay (ELISA, Enzo…) following the manufacturer’s instructions. The Enzo Life Sciences ELISA kit was preferred over other kits because it was showed the highest sensitivity in a previous study^30^. Corticosterone was chosen over cortisol as a stress marker due to its favorable dynamics corresponding more directly to the chronic stress response^19^. The assay was performed in duplicate and plates with final samples after running all the procedures were read on PHERAstar FS, a multi-mode microplate reader (BMG LABTECH Inc., NC, US). Open-access online analysis service (MyAssays Ltd., US) was used for plotting the standard curve and data extrapolation. *Blood corticosterone concentration/corticosterone level* refers to the amount of corticosterone in plasma and is always expressed in nanograms per milliliter.

In the 10-day preparatory experiments, 25-day protocol and 5-day protocol of the supplementary study single blood samples were collected at the end of the session on the first day and every 5 days until the end of the experimental protocol. Furthermore, in the 25-day protocol additional control samples were collected 5 and 30 days after the last head-fixed session and in the 5-day protocol, 5 days afterwards. Also, control experiments for the blood sampling procedure were performed by running 5 pairs of the head-fixed/control animals in the 25-day protocol and withdrawing them from study at different times after a single blood sample performed on day 5, 10, 15, 20 and 25. Moreover, 8 single-housed animals underwent the same preparatory/surgical procedures and the full 25-day protocol without head-fixing and with the same 5-day blood sampling procedure for comparison with the pair-housed animals that were never head-fixed and were used as controls in the 25-day protocol. In addition, in the 5-day preparatory studies, 3 pairs of the head-fixed/control animals were blood sampled 3 times per day for 5 days.

### Behavioral tests

Effects of extended head-fixation were tested in classical stress-associated behavioral paradigms. Behavioral experiments started after the last day of head-fixation in the 25-day protocol and in the 5-day protocol of the supplementary experiments and they were performed on the individual animals in the following order: morning of day 1 – open field test (OFT); afternoon of day 1, 6-hours later – forced swim test (FST); morning of day 2 – elevated plus maze (EPM); day 2/3 overnight – nesting behavior test (NBT) and habituation to two-bottle choice used for the sucrose preference test; day 3/4 – SPT; day 5 – control blood sampling. Mice were tested individually in all the behavioral tests. In OFT and EPM all the surfaces were cleaned with 70 % ethanol in-between running individual animals. In OFT, FST and EPM mice were transferred to the experimental room about 45-60 minutes before the beginning of the test. The same video camera Logitech HD Pro Webcam C920 (Logitech International SA, Switzerland) and the same video tracking analysis software EthoVision XT 13 (Noldus, The Netherlands) was used for all the recordings in OFT, FST and EPM.

In OFT the mouse was placed in the corner of a black plywood box without the top and the bottom (50 × 50 ×40 cm) standing on a light grey base. Every session lasted 10 minutes and was recorded with a video camera positioned above the center of the box. FST was performed in a transparent plastic cylinder filled with water (about 22-25°C). Each animal was placed in the water for 6 minutes and movement/immobility was recorded with a video camera positioned approximately 2 meters in front of the cylinder. After the test the wet mouse was placed on a paper towel in a holding cage warmed up with a heating pad up to 37°C for about 15 minutes to dry. Then it was returned to its home cage. EPM consisted of two open and two closed arms (all arms: 30 × 5 cm) made of black plastic. The floor was made of white plastic for better contrast and it was elevated 50 cm above the ground. Closed arms had 15-cm-high black plastic walls at the sides and at the end. Each animal was placed in the middle of the EPM for 7 minutes and was recorded with a video camera placed above the center of the EPM. The first minute was skipped in the analysis to avoid random choices between the open and the closed arms in the early exploratory phase. NBT was performed following the guidance in the original paper by Deacon ^25^. In short, mice were placed in new cages with 3 grams of untouched nesting material and food pellets on the floor and they were left for the night. In the morning of the next day animals were transferred to new cages and the quality of nests was scored on a scale from 1 to 5 (the worst to the best quality, respectively). SPT was performed using a two-bottle free choice method in which each mouse was presented simultaneously with two bottles (25 ml). Initially, animals were habituated overnight to the two-bottle choice (both bottles were filled with tap water and placed through the top of the cage lid). Then mice were transferred to new cages where one bottle contained tap water and the other 1 % sucrose solution. After 24 hours the position of the bottles was switched to control for a side preference in drinking behavior. On the final day the bottles were weighed, and the sucrose preference index was calculated based on the proportion between the volume of the liquids consumed.

### Locomotion analysis

Head-fixed mice in the MHC apparatus rested their paws on the air-lifted floating container. Thus, every movement of their paws corresponded to an immediate movement of the container that is easily recorded and tracked. Video recordings were collected with a high-definition camera Logitech HD Pro Webcam C920 (Logitech International SA, Switzerland) mounted above the head-fixed apparatus. Video-tracking software EthoVision XT 13 (Noldus, The Netherlands) was used to track the movement of a yellow sticker attached to the outer wall of the floating container. Movement of the floating container was used as an approximate measure for a mouse’s body movements/voluntary running in all the experimental protocols. In the fixed group of the 5-day protocol, in which the container was fixed to an air-table, instead of the yellow sticker, mouse’s body movements were tracked directly (mostly movements of the paws and the tail) using the same tracking software.

In the floating group under the 5-day protocol in the supplementary experiments, the MHC Tracking System (Neurotar Ltd, Finland) was used in addition to the EthoVision software and the Logitech camera. The system consisted of a magnetic board installed inside the air-dispensing base and a rubbery mat with two built-in magnets inserted into the floating container. Air pressure was adjusted because floating container with a rubber mat insert is heavier. Also, it was easier for a mouse to move the container with the insert because it had a much rougher surface providing an easier grip for a mouse. Changing position of the magnets during the head-fixed session was recorded and movement data were extracted with the accompanying software. This setup allowed for detailed analysis of spinning behavior (see Results for definition) as well. With the help of the accompanying software, the MHC container space was divided into two zones: a wall zone (area within 1 cm from the wall towards the middle of the floating container) and a middle zone (the rest of the inner space of the container). The time that the mouse spent in each zone during the head-fixed session was quantified. Moreover, it was possible to color-code their movement. The track maps corresponded to the time when the mouse was in a certain zone. The spectrum of colors from blue to red indicated the time into the head-fixed session, with blue used for the early times and red for the later times.

In the 25-day protocol motor skill refinement was quantified. Namely, transition from more random to organized and controlled movements was quantified with detailed velocity analysis. Temporary changes in velocity called *bouts of activity* were extracted and classified (for raw data, see Supplementary fig. 5a-c). There were 4 bout categories distinguished based on their duration and velocity: long & slow, long & fast, short & slow, short & fast. The cut off value for short-to-long duration was 5 seconds that corresponded to an average time in which mouse was able to move the container for one full lap/circulation. The cut off value for slow-to-fast velocity was 600 cm/min (about 10 cm/sec), because it corresponded to the maximum velocity during the initial phase of learning before animals become efficient in control of the floating container (for raw data, see Supplementary fig. 5d).

### Statistical analysis

Data in the text and on the figures are presented as means ± SEM. Two-tailed paired Student’s t tests were used for comparisons between two groups when conducting single factor analysis. Ordinary or repeated measures one-way ANOVA with Tukey’s or Dunnet’s post hoc multiple comparisons tests were used when one factor was measured at different times for a single group. Two-way repeated measures ANOVA with Tukey’s or Sidak’s post hoc multiple comparisons tests were used when one factor was measured at different times for two groups. Pearson’s correlations and linear regressions were performed to assess the relationship between two independent variables. Data were analyzed with the GraphPad Prism7.04 (GraphPad Software, CA, US).

## Acknowledgements

We thank all the members of the Lovinger and the Alvarez laboratories for their generous discussions and helpful feedback. In particular we thank Dr. Daniel da Silva for the blood sampling technical training, Dr. Miriam Bocarsly for the suggestions on the behavioral paradigms, Dr. Armando Salinas for recommendations on the corticosterone measurements and Dr. Sebastiano Bariselli for critical reading of the manuscript. We also thank Dr. Raouf Kechrid for his helpful suggestions regarding the blood sampling technique and Dr. Veronica Alvarez for her overall scientific support. This work was supported by the Division of Intramural Clinical and Biological Research of the NIAAA (ZIA AA000416).

## Author Contributions Statement

K.J. and D.M.L. conceived the project. K.J. conducted the head-fixed experiments with the help from J.A.K. Moreover, K.J. planned and performed the blood sampling procedure, the behavioral experiments, the blood corticosterone concentration analysis and the locomotion analysis. A.J.K. actively participated in designing behavioral experiments and performed a part of the behavioral analysis. J.O.L. actively participated in designing and performed a part of locomotion analysis. K.J. and D.M.L. wrote the manuscript with input from J.A.K., A.J.K. and J.O.L.

## Additional Information

The authors declare no competing financial interests.

