## Supplementary information for "Stress and behavioral correlates in the head-fixed method"

**Supplementary figure 1. Corticosterone dynamics in the pilot and the control experiments.** (a-i) Blood corticosterone concentration data. (a) 10-day pilot experiment; statistically significant difference between the groups, two-way RM ANOVA: interaction (p = 0.3344), group (p = 0.0003), time (p = 0.0529). Sidak's multiple comparisons between groups: difference at D1 (##p = 0.0152) and D5 (#p = 0.0152), no difference at D10 (p = 0.0538). Tukey's multiple comparisons within groups: in the head-fixed D1 vs. D10 differed significantly (*p = 0.0129), no difference in D1 vs. D5 and D5 vs. D10 (p > 0.19) and in any comparison in the control group (p > 0.4); n = 3 in each group. (b-f) 5-day pilot experiments, 3 blood samples every day (time 0, 60, 120 minutes); n = 2 in the head-fixed and n = 3 in the control group. (b) Percentage change in the corticosterone level during the head-fixed session normalized to the baseline collected at the time 0 minutes. (c) Blood corticosterone concentration at 120 minutes. (d-f) Data for individual animals at the time 0, 60 and 120 minutes. (g, h) Blood sampling control data from individual animal in the 25-day protocol, a comparison between the animals that were blood-sampled several times with the animals sampled only once at a specific time (several vs. single BS subgroups); data collected at day 5, 15 and 25 of the head-fixed protocol; n = 5 in each group. (i) Effects of extended head-fixation protocol on the circadian rhythm of corticosterone; all measurements from the 25-day protocol combined for every blood sampling time; time of the day measured from the beginning of the light cycle (Zeitgeber time, ZT); n = 2 in each group at every time. (j) Control experiments for single housing; a comparison in the corticosterone level between single- and pair-housed non-head-fixed animals, no statistically significant difference between groups but significant effect of time; two-way RM ANOVA: interaction (p = 0.9406); group (p = 0.8661); (p = 0.0313); Tukey's multiple comparisons within groups: no significant difference in any of the comparisons (p > 0.2 in all cases); n = 7 in each group.

**Supplementary figure 2. Complementary data for the behavioral tests.** (a, b) Open-field test: no statistically significant differences between the groups in the number of returns to the central part of the open field box and the latency to the first visit in the center of the open field box; p = 0.7662 and p = 0.6085, respectively. (c, d) Forced-swim test: no statistically significant differences between the groups in the number of the floating events and the number of feces left in the water after the trial; p = 0.7662 and p = 0.6891, respectively. (e-l) Elevated plus maze task: no statistically significant differences between the groups in the number of returns to the center, to the closed and to the open arms (p = 0.6137, p = 0.9418 and p = 0.5669, respectively); the latency to the first visit to the center, to the closed and to the open arms (p = 0.6590, p = 0.3539 and p = 0.6968); the total time spent in the center and in the closed arms (p = 0.6808 and p = 0.8650).

**Supplementary figure 3. Individual differences in the voluntary running in the 25-day habituation protocol.** (a) (b, c, d) Changes in the average movement time per day over 25 days, animals organized in 3 groups, no change, statistically significant increase and statistically significant decrease.

**Supplementary figure 4. Single animal example of data used for analysis of the *bouts of activity*.** (a-d) All data were obtained with a 30-Hz camera and modified for a final analysis to a resolution of seconds to avoid movement artifacts. (a-c) Changes in velocity dynamics during a 120-minute head-fixed session throughout 25-day protocol (data from day 1, day 15 and day 25, respectively). Visible increase in a bout frequency as well as velocity. (d) Data used for distinction between the slow bouts and the fast bouts. Bouts were organized by their velocity in bins with 50 cm/min increments (e.g. 0-50 cm/min.; 50-100 cm/min., etc.). Histogram representing all bouts from the first 5 days (in blue) and the last 5 days (in red) of the protocol and depicting their total number. Distribution of different velocities was much broader at the end of the 25-day protocol. The cut off value for the slow and the fast bouts was 600 cm/min (about 10 cm/sec) because it corresponded to the maximum velocity during the initial phase of learning before animals become efficient in the control of the floating container.

**Supplementary figure 5. Corticosterone dynamics of the head-fixed animals in the 5-day head-fixed protocol followed by stress-associated behavioral tests.** (a-f) 5-day head-fixed protocol. (a-c) Blood corticosterone concentration; n = 4 in each group. (a) Statistically significant drop in the corticosterone level in both groups at D10; two-way RM ANOVA; interaction (p = 0.7163); group (p = 0.3470); time (p = 0.0001); Tukey's multiple comparisons within groups: D1 vs. D10 (**p < 0.01 in both groups), D5 vs. D10 (**p = 0.0030 in the floating and *p = 0.0109 in the fixed group), D1 vs. D5 (p > 0.8 in both groups).(b, c) Individual animal data for the floating and the fixed group. (d-f) Movement time analysis; n = 3 in each group. (d) Significantly shorter movement time in the fixed group but no change throughout training days, two-way RM ANOVA: interaction (p = 0.1950), group (*p = 0.0303), and time (p = 0.1306). (e) Percentage change in the movement time normalized to D1. (f-q) Data from all the behavioral tests; dashed line corresponds to the head-fixed group from the 25-day protocol; paired t-test in all comparisons; n=4 in each group. (f, g, h) Open field test: the total distance traveled, the time in the center and the latency to the first open arm (**p < 0.05), respectively. (i, j, k) Forced-swim test: the latency to the first floating event, the total floating time and the number of the floating events. (l, m, n) Elevated plus maze: the total distance traveled, the time in the open arms and the number of attempts to the open arms (*p = 0.0349). (o) Nesting behavior: the nesting quality score. (p, q) Sucrose preference test: the sucrose preference score and the total volume consumed in the floating group (**p = 0.0032), bottom dashed line = water, and top = sugar.

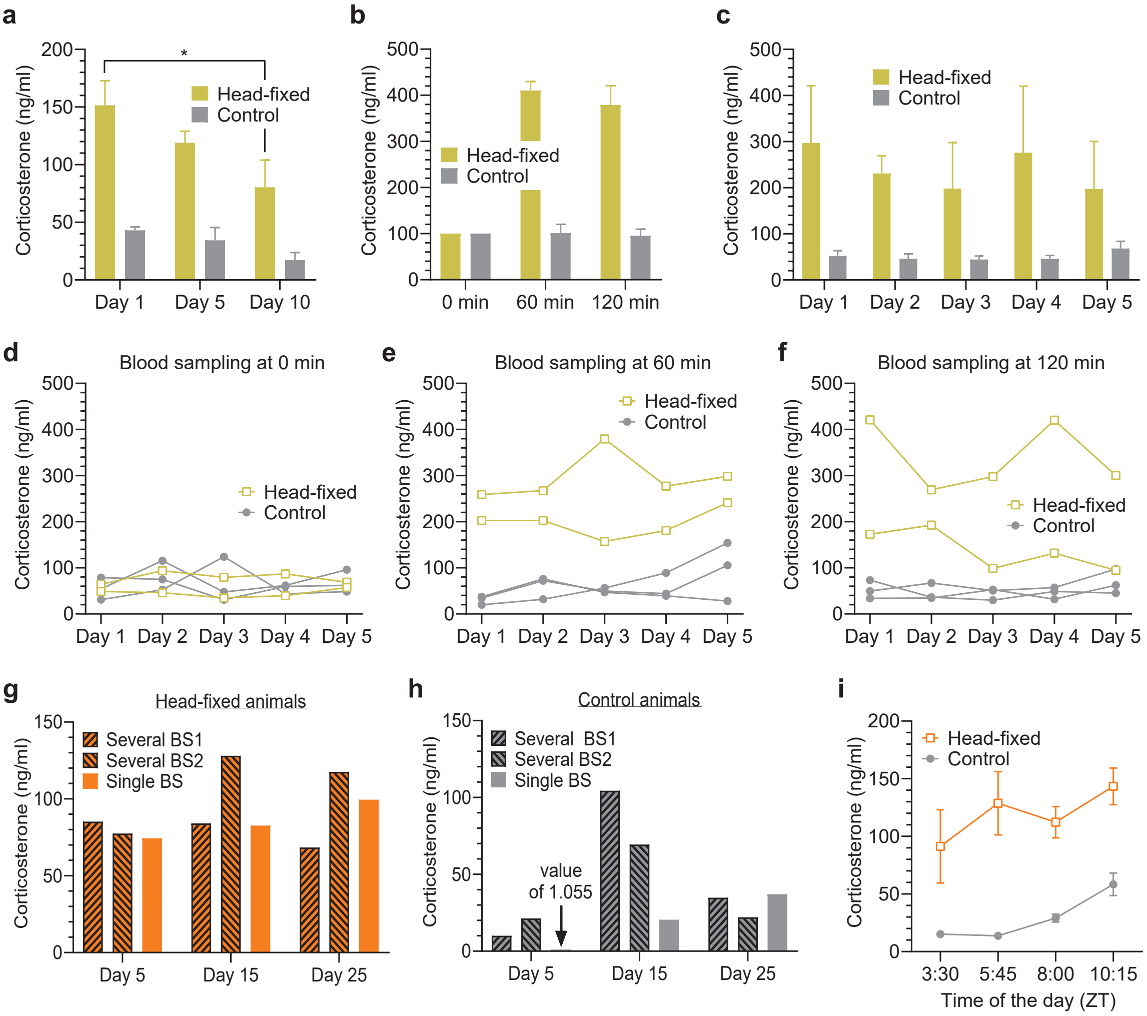


**Supplementary figure 1 (Fig. S1)**

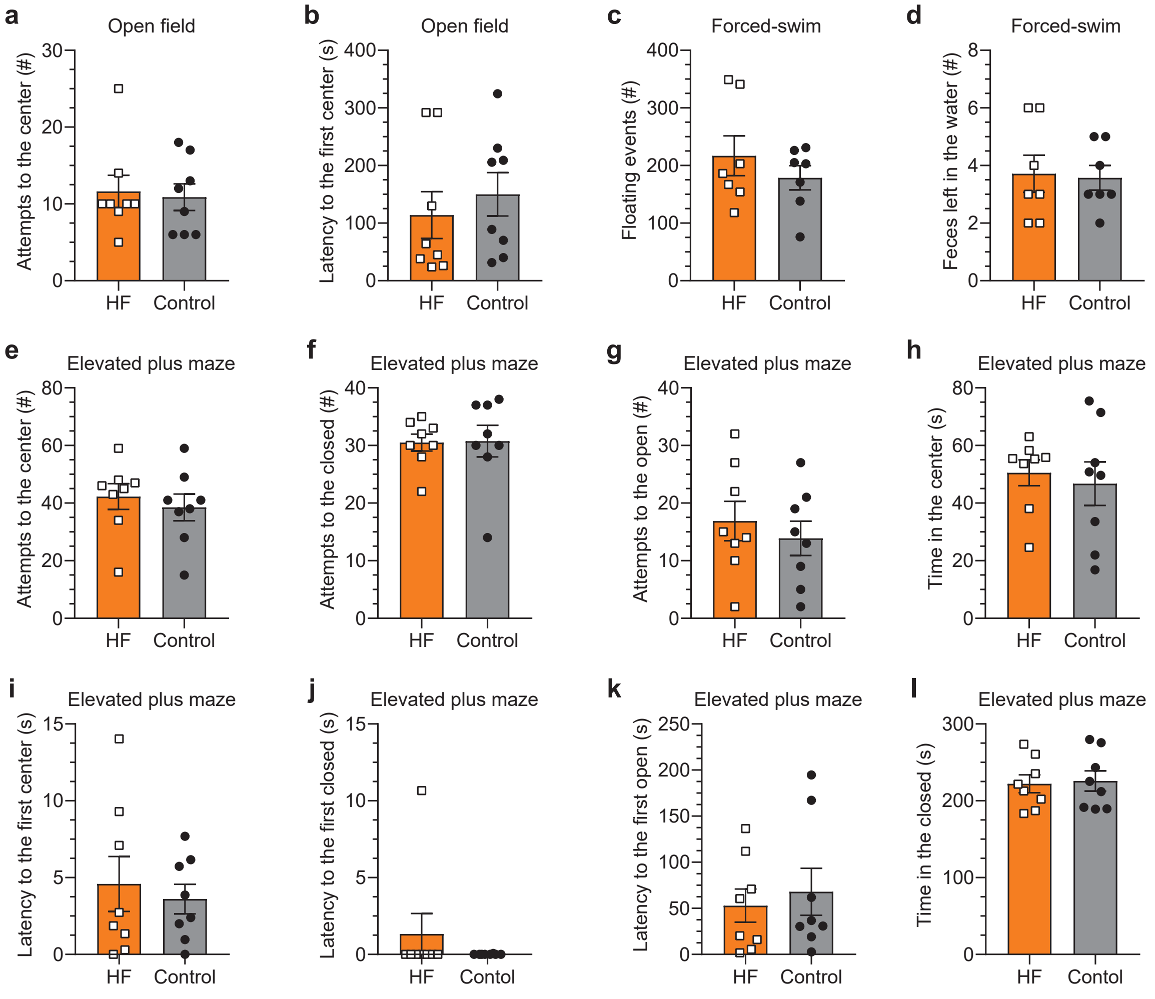


**Supplementary figure 2 (Fig. S2)**

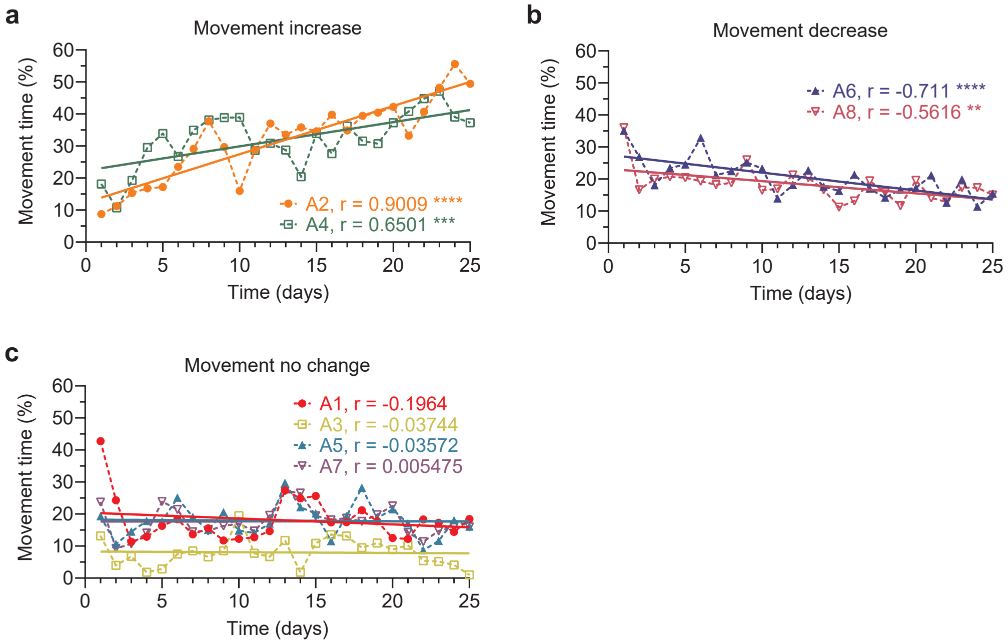


**Supplementary figure 3 (Fig. S3)**

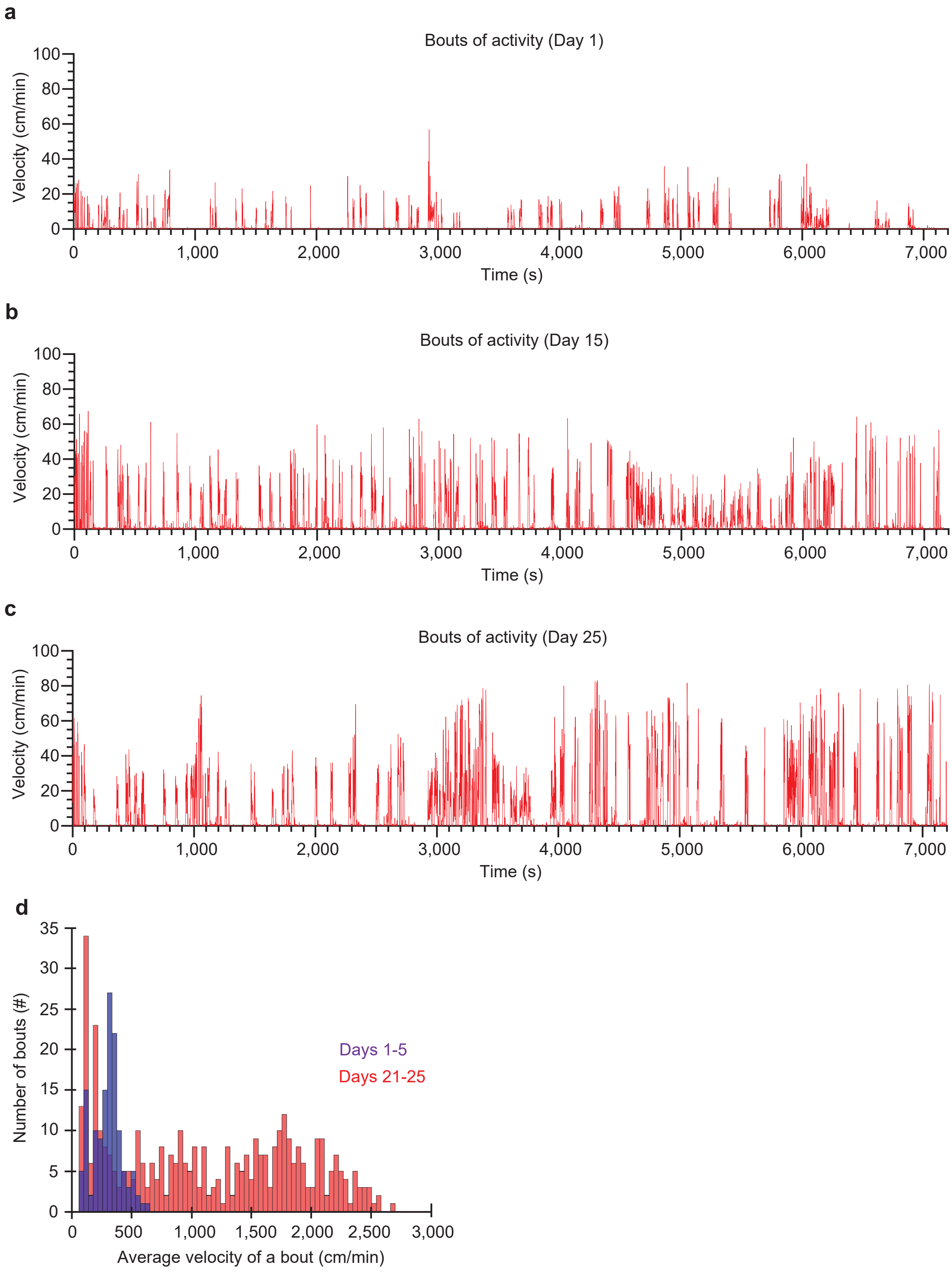


**Supplementary figure 4 (Fig. S4)**

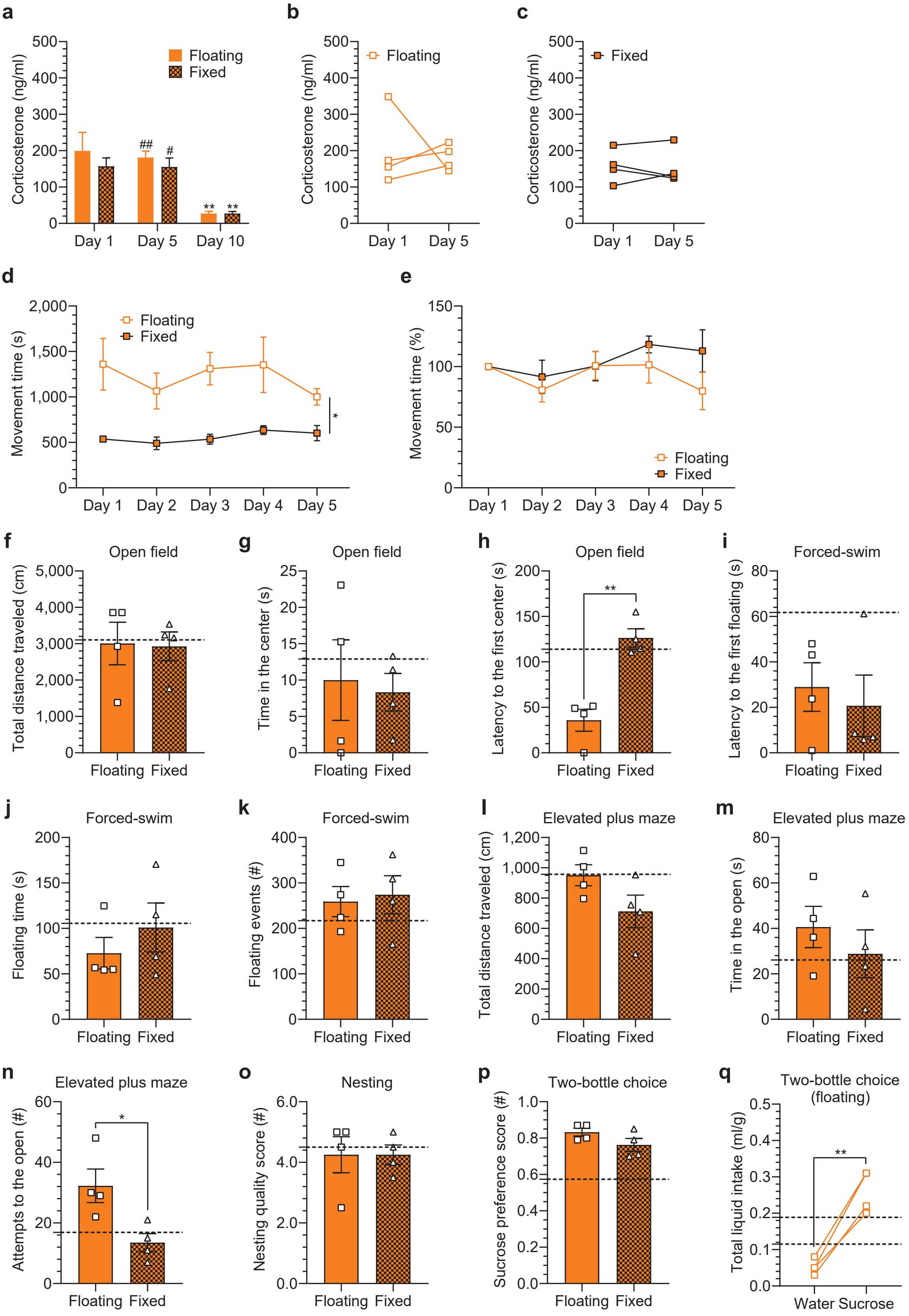


**Supplementary figure 5 (Fig. S5)**
